# A Closer Look at the White Pine Cone Beetle

**DOI:** 10.64898/2026.05.21.726841

**Authors:** Apolline Maurin, Audrey-Anne Durand, Philippe Constant, Claude Guertin

## Abstract

*Conophthorus coniperda* (Coleoptera: Curculionidae; Schwarz), known as the white pine cone beetle, is a pest in white pine seed orchards. The insect was primarily studied in the United States of America during the late 1960’s to 1992. No contemporary studies have been performed since, despite the devastating damage the beetle causes to white pine seed orchards, and the seeds they produce for reforestation purposes. To help future research on potential biological control, this work revisits the ontogeny of the white pine cone beetle. The biology of the insect was studied over two years in 2022 and 2023 in a seed orchard (Québec, Canada). Observations were complemented with data collected from the same orchard and other sites in 2009 and in 2012. Except in 2022, the emergence of the insect occurred around 53.6 ± 1.98 °C.d above a threshold of 6.5 °C. A shorter developmental cycle was observed compared with the ones described in 1965 and 1976. The relationship between cone size and the number of insects per cone was statistically significant but explained only a small proportion of the variance and showed high variability. These results will help improve survey timing and sampling strategies for this species.

## Introduction

The white pine cone beetle (WPCB), *Conophthorus coniperda* (Schwarz; Curculionidae: Scolytinae), is a small beetle (2-4 mm) that attacks cones of white pines, *Pinus strobus* (Linnaeus), and feeds on their seeds. White pine cones follow a two-years developmental cycle (Krugman & Jenkinson, 1974). After cone fertilisation in spring or early summer, cones begin to develop and will mature over the following year. The WPCB primarily targets second-year cones (Godwin & Odell, 1965). The population density of *C. coniperda* is influenced by the annual fluctuation in seed and cone production (Turgeon *et al*., 1994).

In spring, adult females emerge from cones where they have been overwintering, and initiate attacks on new cones at the junction between the cone and the peduncle (Godwin & Odell, 1965). After a feeding period, female beetles release a sexual pheromone, pityol, to attract males for mating. This pheromone may be released alone, but attracts more males in association with monoterpene found in cone (Birgersson *et al*., 1995, Brauner & de Groot, 2006). After mating, the female lays eggs in niches along the side of gallery dug in the cone axis. Eggs hatch in about a week, followed by two larval instars, each lasting about two weeks, and pupation. Callow adults appear 7–10 days after pupation, eventually developing into fully sclerotized, ready to emerge, adults (Godwin & Odell, 1965). A single female will attack two to four cones during spring, either to feed or to breed (Godwin & Odell, 1965). Attacked cones are systematically aborted by the tree, resulting in seed mortality of over 80% (Trudel *et al*., 2004, Guertin & Trudel, 2006).

Even though the WPCB leaves the rest of the tree undamaged, it is often seedless (Trudel *et al*., 2004, Guertin & Trudel, 2006). It is therefore a significant pest in seed orchards, where white pine trees selected for superior form or growth produce seeds in abundance for reforestation purposes (MRNF, 2019, MRNF, 2024). Indeed, white pine has been part of the reforestation program since the 1980s due to the overharvesting of the tree during the 19^th^ century and its patrimonial and economical value in Canada.

Various controls methods have been considered (e.g. pesticides, prescribed burning and pheromone-mediated mating disruption) to cope with this devastating pest, without success (Godwin & Odell, 1965, Trudel *et al*., 2004, Guertin & Trudel, 2006, Guertin & Trudel, 2007). Disruption of insect-microbiome symbiosis is envisioned as a promising biocontrol strategy (Lv *et al*., 2024). Insects are closely related to microorganisms (bacteria, fungi, archaea, viruses) (Popa *et al*., 2012). They play crucial roles in the protection, alimentation and establishment of their insect host (Coon *et al*., 2020, Li *et al*., 2021, Diehl *et al*., 2022). Hence, disruption of the insect’s microbiome is expected to influence the fitness of insects. However, the role of microorganisms on the WPCB fitness and the variation of this beetle’s microbiota in response to environmental cues are critical knowledge gaps to bridge before the implementation of insect-microbiome symbiosis centric biocontrol. To successfully apprehend the influence of microorganisms on the fitness of WPCB, it requires to actualise the knowledge on the biology of this beetle. A better understanding of its ontogeny, phenology and behaviour will influence the experimental design and the period of collection.

Therefore, key biological traits of the WPCB in Québec (Canada) have been investigated and compared with published studies and/or archived data collected by our research group throughout the years.

## Materials and Methods

### Insect collection

The biology of the WPCB was monitored over 2 years, in 2022 and 2023, in the white pine seed orchard of Verchère, located in Saint-Amable (Québec, Canada, Lat. 45.677000, Long. -73.330000) with the permission of the Ministère des Ressources Naturelles et des Forêts (MRNF). The orchard comprises around 3 758 trees distributed over 35 815 m^2^. Cones were collected on the ground from 18 distinct locations of the orchard, selected randomly (Supplemental Materials 1). The collection was done in a 5 meters radius around the selected tree, which comprises up to 6 additional trees. For each, up to 12 cones were collected weekly, bimonthly or monthly from April to November in 2022, in February 2023 and from April to July in 2023 (Supplemental Materials 1). Collected cones were preserved in a cooler during transportation and then stored at 4 °C until treatment. All samples were processed within 24 hours of collection. Six parameters were measured or assessed in this study: (i) adult emergence; (ii) feeding before mating; (iii) mating; (iv) oviposition; (v) life stages; (vi) behaviour (movement within cone and emergence).

### Insect biology monitoring

To predict the emergence period of the WPCB, heat accumulation was monitored using the cumulative degree-day model calculated with the DegDay software version 1.01 of Richard L. Snyder (Snyder, 2005) using the Sine Wave methods (Allen, 1976) for a threshold temperature (T) of 6.5 °C (Guertin, 2014). This threshold refers to the lower developmental temperature threshold. Meteorological data were collected from Agrométéo Québec (Mesonet, 2025), for Calixa-Lavalée (Québec, Canada, Lat. 45.72935, Long. -73.25378), located at about nine km from the orchard. Results comprised data collected in 2022-2023 and in 2012 (archived data) from the same orchard. Additionally, adult emergence and female feeding period were assessed by observing the presence of young and new cones on the orchard floor, which should contained females only. Newly attacked cone was easily distinguishable thanks to the size, colour and its structural integrity. The mating period was determined using three pheromone traps (Japanese Beetle Trap, Great Lakes IPM™, Michigan) containing pityol (*trans*-pityol Bubble, Solida, Québec) in the trees canopy. Traps were monitored weekly, starting from the first appearance of newly attacked cones and continuing until no males were absent for four consecutive weekly checks. The oviposition period was defined as the time span between the first and last egg found in cones. Then, the succession of developmental stages was registered, and specimens from each stage were photographed under the SterREo Discovery V20 binocular magnifier using the AxioCam Icc 3 (Carl Zeiss MicroImaging, Gottingen, Germany).

### Influence of cone size on the number of beetles per cone

We assessed the potential relationship between the number of insects and the cone length. To this end, data collected in 2009 by our team (archived data) and in 2022 were compiled. Cones from 2023 were not included as they were only used to assess the biology of the WPCB. As a result, 409 cones (in 2009) and 666 cones (in 2022) were measured with a digital calliper from the base of the peduncle to the tip of the cones. Cones were then opened, and insects were counted (Supplemental Materials 1). In 2022, cones were collected exclusively from the Verchère orchard, while in 2009, they were collected in multiple orchards: Cleveland (Lat. 45.6809, Long. -71.9969; 49 cones), Verchère (101 cones), Aubin-de-Lisle (Lat. 46.187134, Long. -70.639931; 50 cones), Cap-Tourmente (Lat. 47.088526, Long. -70.790472; 71 cones), Huddersfield Lat. 45.921258, Long. -76.737115; 138 cones).

As count data were overdispersed relative to Poisson and zero-inflated, a generalised linear model using a negative binomial was used to assess the relation between the cone length (fixed effects) and the number of insects in cones. Analyses were performed using R Statistical Software version 4.3.2. (R Core Team, 2024), RStudio software (Posit team, 2024) and the MASS package version 7.3-61 (Venables & Ripley, 2002). To ensure the model fit our data and significantly explained the variance, the McFadden pseudo r2 was calculated using the pscl package version 1.5.9 (Jackman, 2024). A McFadden pseudo r2 between 0.2 and 0.4 is often considered indicative of good fit. To test the influence of year of collection (fixed effects) and the localisation (i.e. different seed orchards ; Random effects) on the number of beetle per cone, a generalised linear mixed model was used with the glmmTMB packages version 1.1.10 (Brooks et al., 2017). To avoid imbalance, we only compared Verchère 2009 vs Verchère 2022 for year effect. Figures were made using ggplot2 version 4.0.0 (Wickham, 2016). Raw data can be found in Supplemental Materials 1.

## Results

### Emergence

In 2022 and 2023, the start of female emergence occurred 26 days apart, with 2022 showing a 134.4□°C·d higher (T = 6.5□°C; Fig. 1). Similarly, 21 days separate the emergence of the female in 2023 and 2012. However, both emergences happened at 52.2 °C·d. Overall, regardless of the accumulated degree day, the WPCB emerged between the end of march and May.

**Figure 1:**
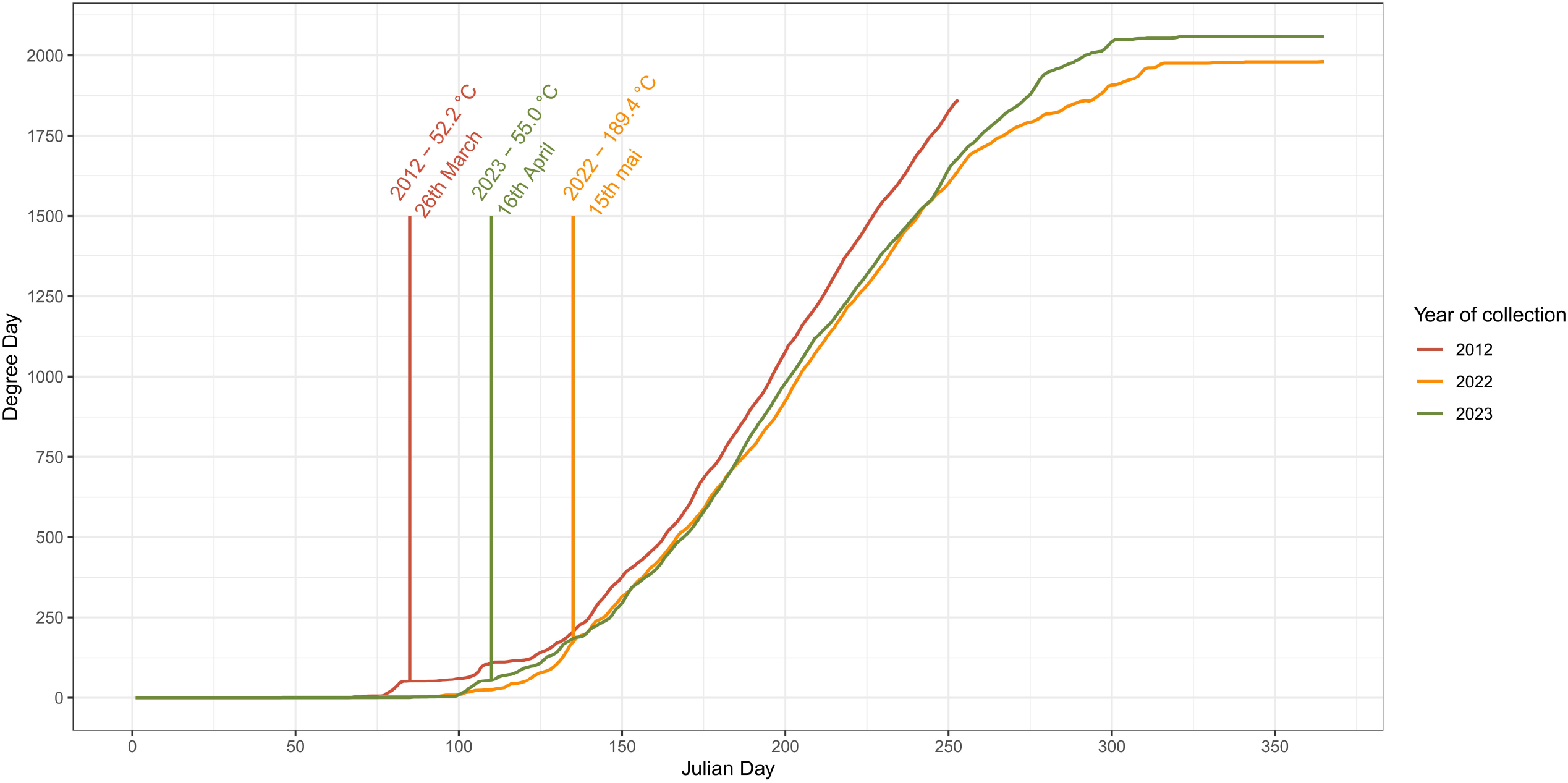
Period of emergence depending on the degree day accumulation above 6.5 °C and the year of collection. The estimated date of emergence is represented by a coloured line. For each year, the heat accumulation and the date of estimated emergence is described.

### Ontogeny and phenology

Males were capture in traps approximately 15 days after emergence and female feeding (Supplemental Materials 1). Before oviposition was observed, we often found several male and/or females in a single cone. The first eggs were collected seven to ten days after first male captures and could be found for two weeks, before larvae appeared in cones (Fig. 2). It is worth noting that oviposition may have started earlier, but cones were still attached at the trees canopy. Their collection was compromised by logistical constraints imposed by the height of the trees in the orchard. There were two larval stages that were mainly found in June but lasted until end of July. The two larval stages could be distinguished by their distinct coloration and by the structural fragility of the second instar. The first instar typically develops a brownish coloration due to active feeding, whereas the second instar remains predominantly pearly white. Pupae followed from the end of June until the end of July, with a peak during the first two weeks of July. Then, callow adults appeared for approximately three weeks, starting from mid-July. Fully formed adults were finally found at the end of July.

**Figure 2:**
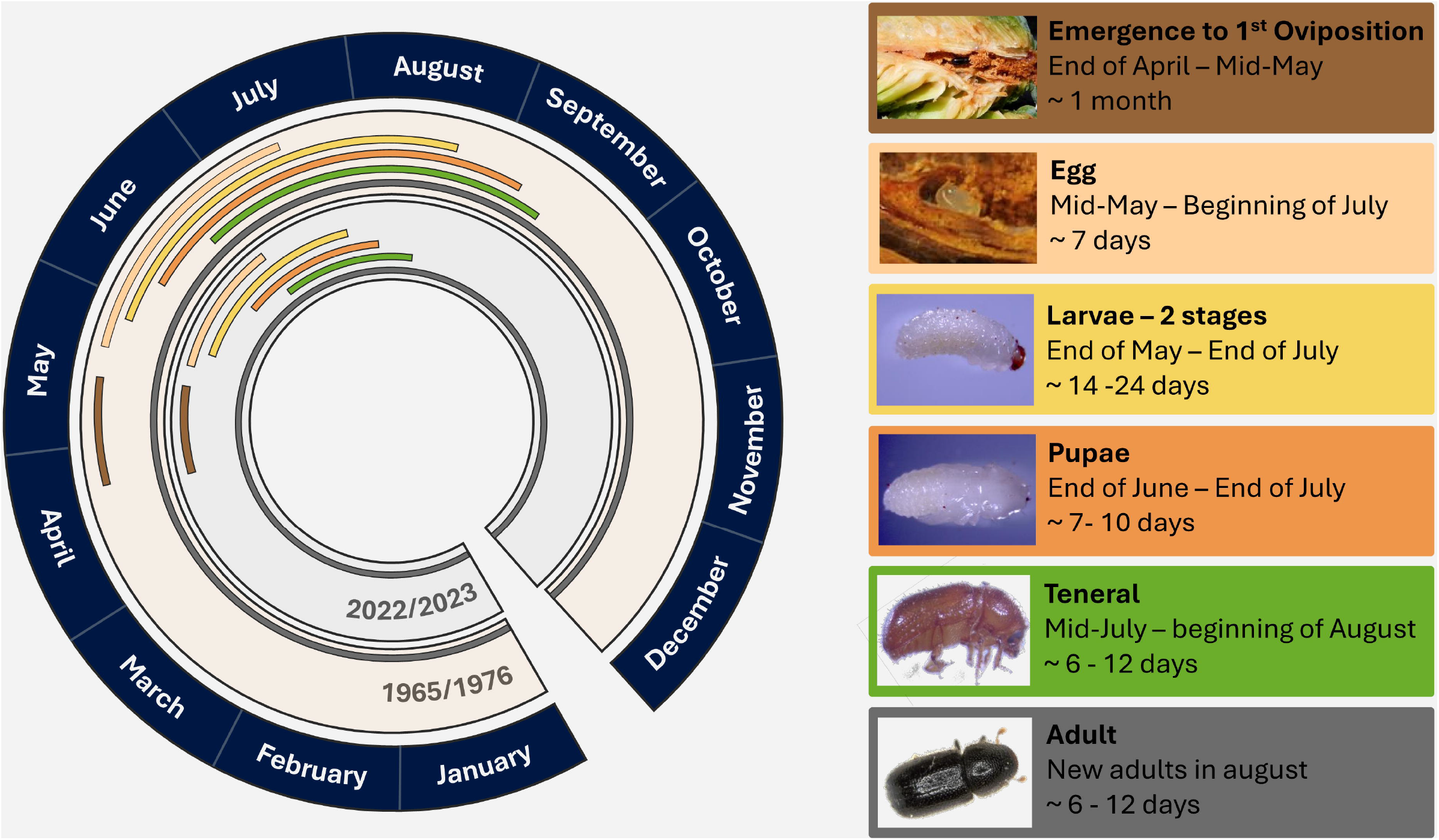
Comparison of the biology of the white pine cone beetle between 1965/1976 (Godwin & Odell, 1965, Morgan & Mailu, 1976) and 2022/2023. (Photos: Apolline Maurin and Guertin’s Lab archives)

### Behaviour

Insects migrate to the peduncles of cones once they become adult. From August, the majority of WPCBs were found digging in the peduncle of cones from inside or outside the cones. Hence, beetles found in the peduncle were not necessarily from the associated cone. This behaviour coincided with the observed occurrence of Lepidoptera larvae from the end of July in most cones.

WPCB leave the brood cones to explore new ones when they reach maturity and before they overwinter. From September, an average of 50 % of the attacked cones on the ground were empty, compared with 19.3 % in July and August (Supplemental Figure 1).

### Cone size influence on beetle numbers per cone

The number of beetles varies greatly from one cone to another. Although typically zero to seven insects were found (in 2009 and 2022), we collected up to 21 insects in a single cone (Fig. 3). In addition, the number of specimens found in cones varies greatly depending on the period of collection (Supplemental Figure 1). It is also interesting to note that the number of insects and its variation over time is similar to the one observed by Godwin & Odell (1965) in Connecticut. Moreover, even if it only explains a small proportion of the variation, cone size is correlated with the number of insects in cones (p_2009_ = 6.73 × 10^−04^ 0.000673, McFadden pseudo r^*2*^_2009_ = 0.08; p_*2022*_ *=* 6.75 × 10^−07^, McFadden pseudo r^2^_2022_ = 6.58 × 10^−03^; Fig. 4). However, the low pseudo r^2^ in the model indicates that other factors than cone size might influence insect density. Notably, the strength of this relation is likely influenced by the period of collection and is representative of the development stage of cones when attacks occurred. On another note, when comparing 2009 and 2022, slopes differ slightly (sd = 0.08; Fig. 4), while they almost do not with the inter-site model in 2009 (sd = 0.02; Supplemental Figure 2). This suggest that the relationship between cone size and the number of insects might have an interannual variation superior to the spatial variation, at the scale of the examined orchards.

**Figure 3:**
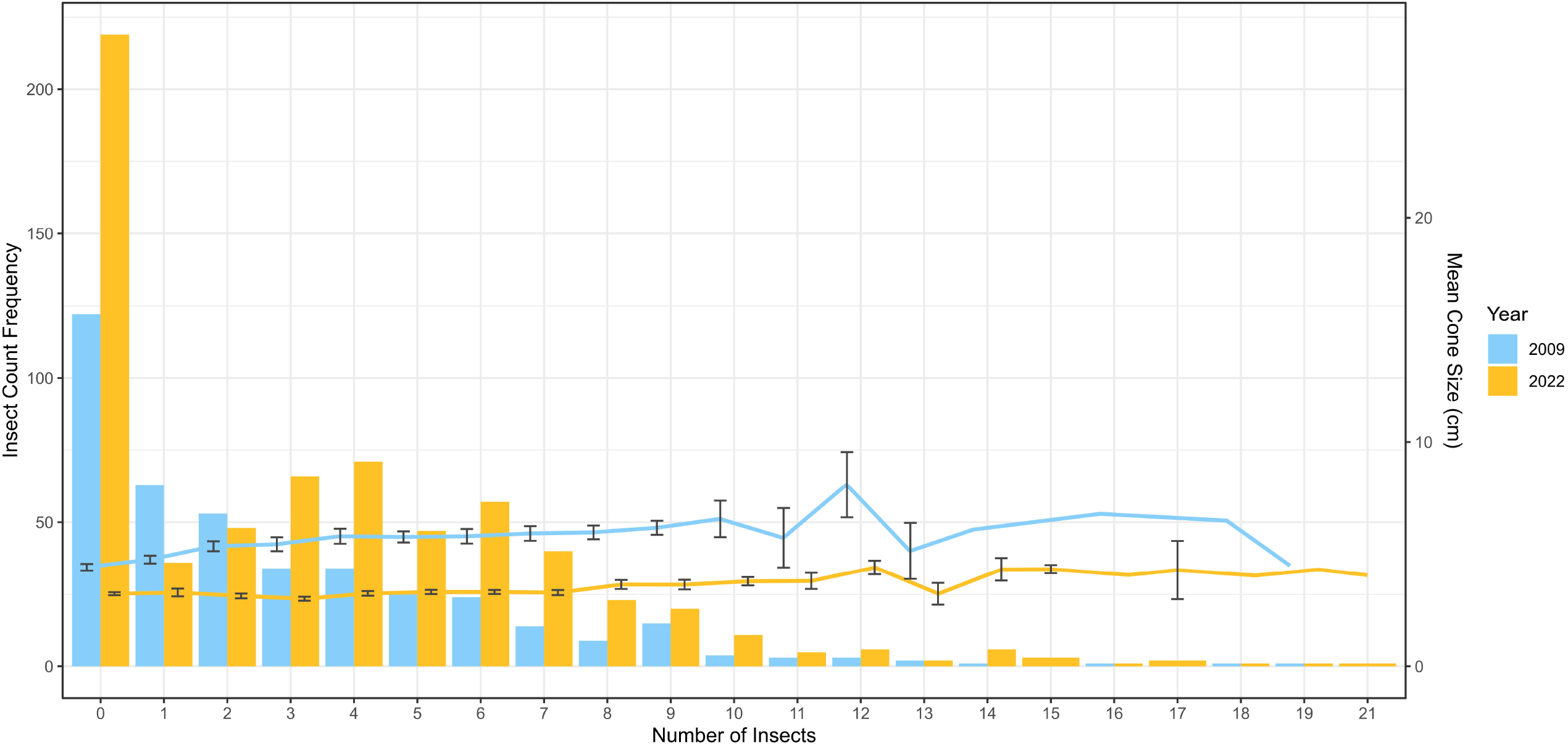
Insect count frequency and mean cone size by year. Histograms represent the insect count frequencies, the curve the mean cone size and the error bars represent the standard deviation.

**Figure 4:**
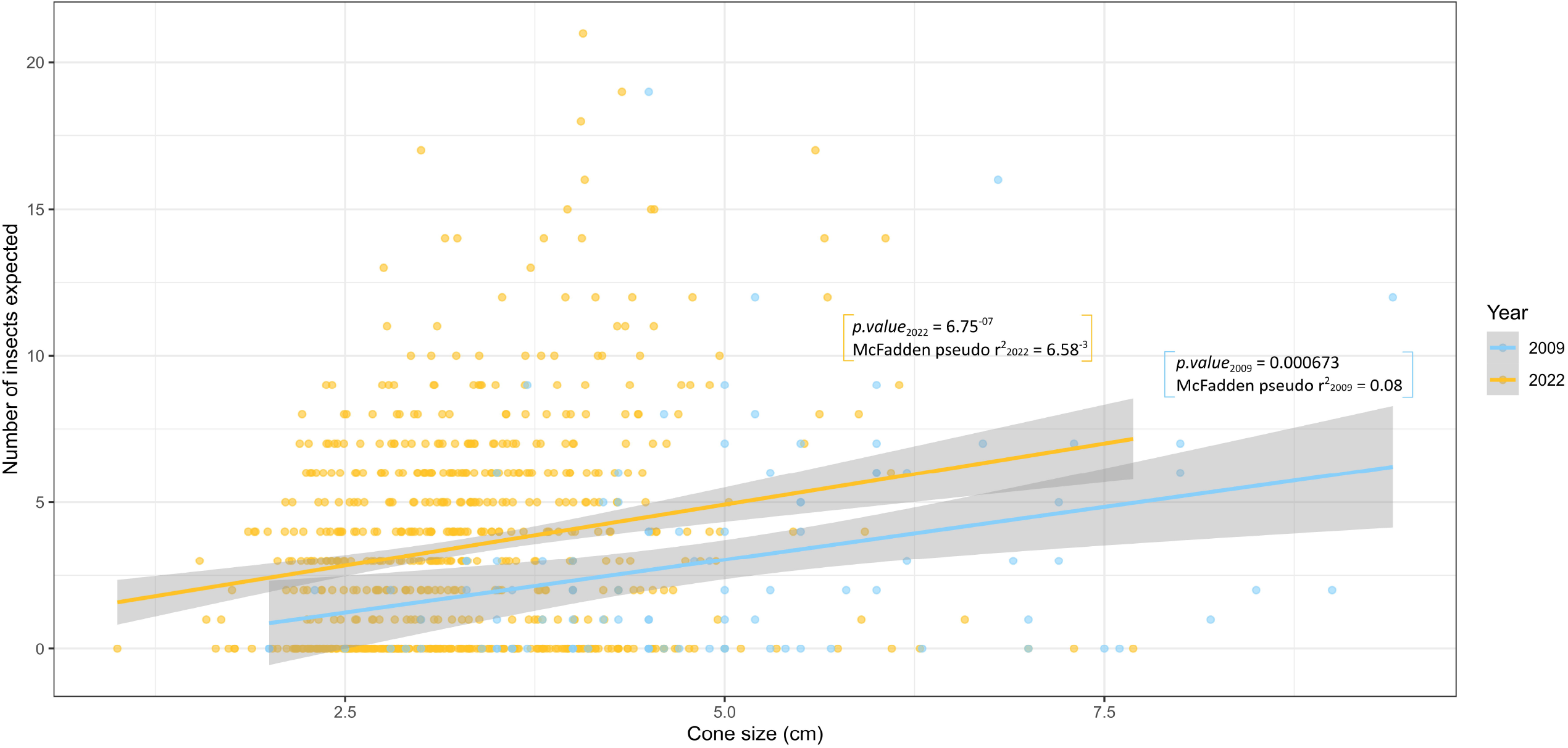
Number of insects expected with cone size in 2009 and 2022 using a generalised linear model.† In memory of Claude Guertin, deceased in July 2023

## Discussion

The WPCB is a significant pest of seed orchards. Despite its impact, the most recent published studies date back to 2005. While the species’ biology has been previously described, ongoing environmental changes, and the known sensitivity of insects to such changes, raise the question of whether the WPCB’s phenology has changed. Additionally, its cryptic behaviour prompts further investigation into whether cone size influences the number of beetles developing within. In this study, we monitored WPCB populations over a two-year period, tracking life stages, the number of insects per cone, and cone dimensions. These findings provide updated insights into the species’ biology and will support the development of effective biological control strategies.

The beginning of female emergence varied from one year to another but shared the necessity of a certain heat accumulation. The difference between 2022 and 2023 may be explained by the slower heat accumulation and persistently low temperatures in 2022, which remained below 20□°C until May 16th. Moreover, despite variation in emergence dates in 2012 and 2023, the average degree-day threshold for emergence remained consistent, occurring around 53.6 □±□1.98 °C.d (T = 6.5□°C; Fig. 1). These observations support the hypothesis of anomalous meteorological conditions in 2022. Regardless of the year of collection, a succession of several days at 20°C or higher appeared essential for the WPCB to emerge, indicating that a minimum ambient temperature might be essential for its emergence. These results are in line with previous studies conducted in laboratory condition. Notably, Henson (1962) worked on the influence of humidity, light and temperature on the WPCB and expected insect to flight under clear and warm circumstances. Especially, beetle seems to require a certain heat accumulation to emerge (Henson, 1964).

The developmental period of contemporary WPCB populations is similar to that reported in the 1960s and 1970s, albeit shorter. Unlike Godwin & Odell (1965) and Morgan & Mailu (1976), the last immature stages were found early August, reflecting a shortening of the developmental period. Otherwise, the development cycle did not differ much from previous studies. The main difference might be the time-lapse where each stage is found. However, our observations from a single site are insufficient to determine whether this difference is due to the location of the studies (Québec vs Connecticut and Wisconsin) or the influence of climate change. In this regard, it is of interest to note that the duration of the WPCB’s development cycle varies from one year to another because of the weather conditions. Indeed, insects are ectothermic and thus highly susceptible to the temperature of their environment (Paaijmans *et al*., 2013, Overgaard & MacMillan, 2017). As a result, the insect’s ontogeny could also be subjected to temperature and the threshold defined for the degree day model might fluctuates according to the stage considered (Kingsolver & Buckley, 2020). A given stage can also be affected by the carry-over effect, i.e. the indirect influence of environmental conditions experienced by earlier stages (Poitou *et al*., 2022). Therefore, the development period of each stage may vary according to the weather and the optimal developmental temperature of each stage of the WPCB (Jaramillo *et al*., 2006, Forrest, 2016). Furthermore, several studies described changes in the phenology of insect and in voltinism in response to climate change (Mitton & Ferrenberg, 2012, Forrest, 2016, Poitou *et al*., 2022). Similarly, climate warming may shift the phenology of *C. coniperda* in a way that enables a partial or full second generation. However, this univoltine species depends on cone availability (Turgeon *et al*., 1994) and although overwintering is not a true obligatory diapause, it coincides with cone maturation. In addition, a single generation typically destroys nearly all available cones, limiting resources for subsequent broods (Trudel *et al*., 2004, Guertin & Trudel, 2006). Since the WPCB is largely host-specific (Henson, 1961), bivoltinism would require exceptionally high cone production or the presence of an alternative host. Such conditions make a second generation plausible, but unlikely.

WPCB begin to co-occur with lepidopteran larvae in August, a period that coincides with their migration into the peduncle. While some of these larvae are known to coexist with the WPCB, others appear to be predators (Godwin & Odell, 1965). Thus, beetle migration from the cone to the peduncle might be a biological response to cohabitation and/or predation, allowing them to protect themselves. The level of cohabitation could be influenced by the intensity of the attacks conducted in the orchards by the WPCB population. Inherently, the level of cohabitation will depend on the size of the beetle population and on the yearly cone production. Another hypothesis is that the peduncle provides a better location for overwintering than inside the cones. Then, the WPCB will migrate into the peduncle regardless of the presence of other insect in the cones. In fact, Godwin & Odell (1965) noted that *C. coniperda* was consistently found overwintering in the peduncle, regardless of whether it was in its brood cone, a newly attacked cone in trees or a previously attacked cone on the ground. To test these hypotheses, the migratory behaviour of *C. coniperda* should be evaluated while considering different site, over several years and with assessing the presence/absence of Lepidoptera in the orchard or without any other insect in the laboratory.

Newly formed adults leave their brood cones in autumn. Similar observations have been described in the work of Godwin & Odell (1965). The benefit of such behaviour for the beetle is not clear. However, during autumn, cones easily lose their peduncle due to humidity. If it is the preferred location of the WPCB for overwintering and/or for protection, it could be reason enough for the insect to change cones. In this sense, the kind of attacked cones might depend on what is available at the time of the autumnal emergence, but also the meteorological conditions.

Finally, the number of beetles found in cones is slightly influenced by the cone length, but the model presented a low explanatory power. These results are in line with the co-dependency of this species to the cone and seed production of white pines (Turgeon *et al*., 1994). It also highlight that the two-years development of cones is influenced by abiotic parameters which influence the morphology and the abundance of the second-year cones (Krugman & Jenkinson, 1974). Similarly, cone maggots have been reported to prefer larger cones for oviposition, as these contain more seeds and thus provide a better food source (Fidgen *et al*., 1998).

## Conclusion

In conclusion, the biology of the WPCB has remained consistent, with an identical ontogeny, and similar time of emergence and oviposition. However, the time-lapse of these events seems to be shorter than it was in 1965 and 1976. Considering the emergence of *C. coniperda*, the combination of the degree day model and the ambient temperature should be used for sampling, particularly the number of days with temperatures above 20ºC. Finally, to facilitate future sampling, we encourage further studies on the link between cone size, number of insect and the biology of the insect and cones.

## Acknowledgements

The authors would like to thank Narin Srei, Rose Ragot and Marie Bonduelle (Institut national de la recherche scientifique) for their technical assistance for the field sampling and decortication of cones. We would also like to thank Pierre-Luc Chagnon (Agriculture and Agri-Food Canada) for his insights on the generalised linear model.

## Conflict of interest

The author(s) declare none

## Notes

### Competing Interest Statement

The authors have declared no competing interest.

