## Supplementary figures and images for "A Closer Look at the White Pine Cone Beetle"

### Supplemental Figure 1

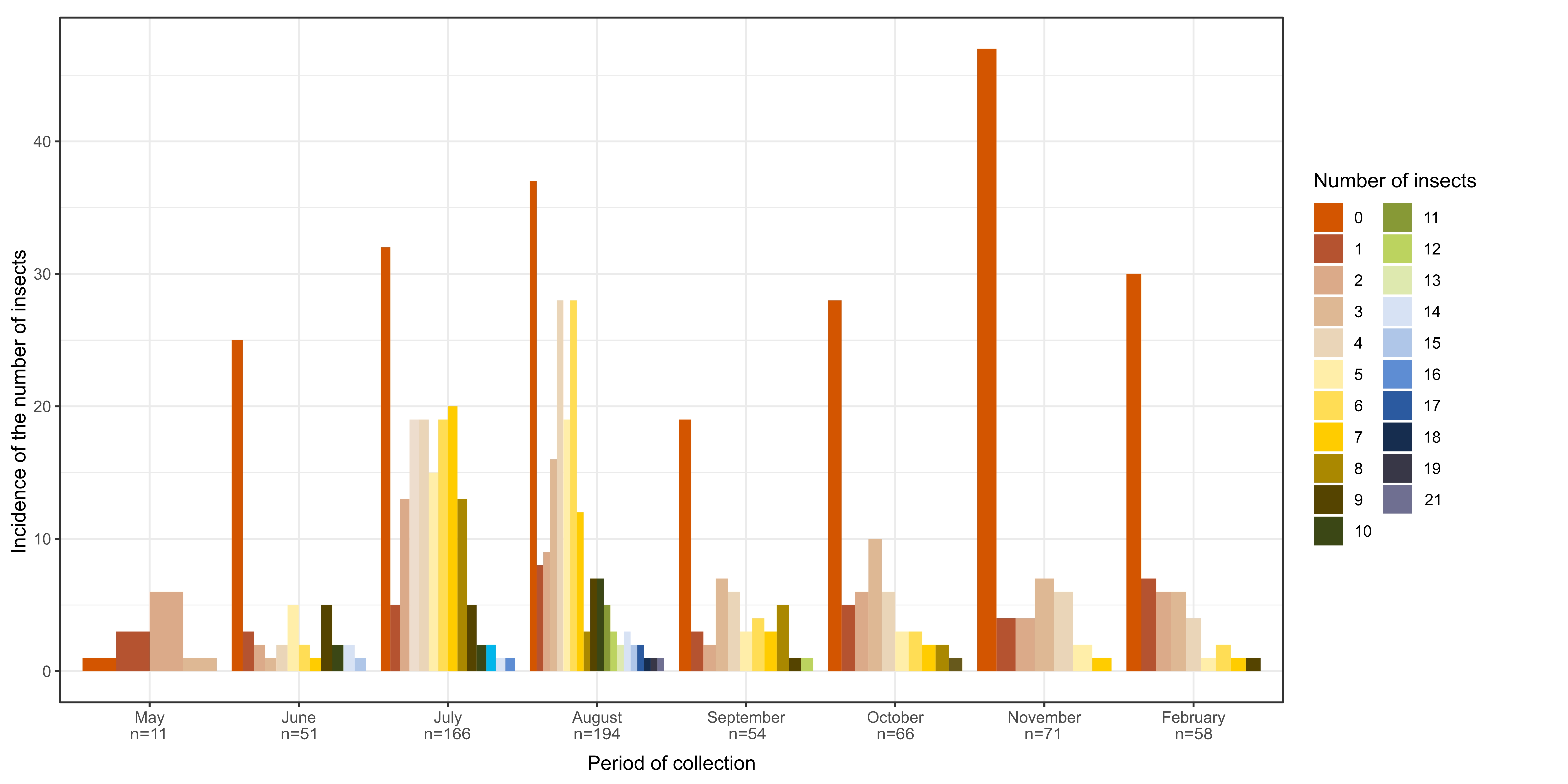

### Supplemental Figure 2

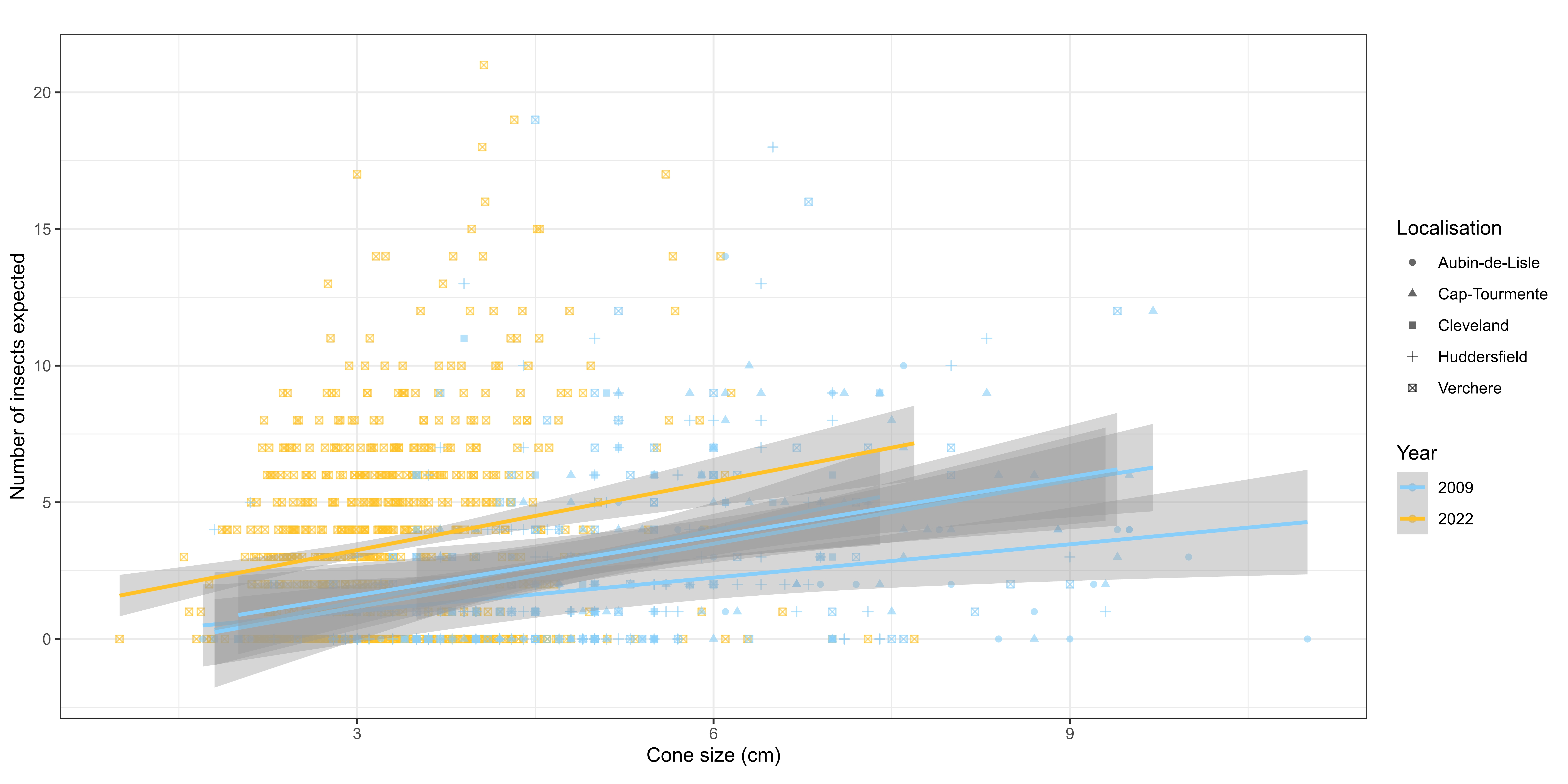
